# Levosimendan inhibits HIV-1 infection in myeloid cells in the RIOK1-dependent manner

**DOI:** 10.64898/2026.04.08.717218

**Authors:** Jinshan He, Jie Ma, Youngmin Park, Dawei Zhou, Xu Wang, Guillaume N Fiches, Kateepe ASN Shanaka, Thurbu T Lepcha, Yan Liu, Suha Eleya, Netty G Santoso, Wen-Zhe Ho, Jian Zhu

## Abstract

Despite of the highly potent antiretroviral therapies, HIV-1 establishes persistent infection and causes chronic inflammation in AIDS patients. Beyond CD4+ T cells, HIV-1 infects myeloid cells, including circulating monocytes and tissue-resident macrophages, and integrates with host genomes to form stable viral reservoirs. To achieve a functional HIV cure, latency-promoting agents (LPAs) have been developed for the “block-and-lock” strategy to reinforce deep HIV-1 latency and permanently silence proviruses. However, most LPAs have been tested mainly in CD4^+^ T cells, and their efficacy in myeloid cells remains unclear. In this study, we reported that levosimendan (LSM), a drug approved for clinic use to treat heart failures, is able to inhibit HIV lytic infection and reactivation in myeloid cells. LSM blocked viral lytic reactivation in HIV-1 latently infected monocytic cells (TH89GFP, U1) and microglial cells (HC69). LSM also inhibited HIV infection in human induced pluripotent stem cell (iPSC) derived microglia (iMG), primary human resident liver macrophages (Kupffer cells) as well as human monocyte-derived macrophages (MDMs). Furthermore, we demonstrated that overexpression of a predicted drug target of LSM, the conserved serine/threonine kinase RIOK1 (RIO kinase 1), overcomes LSM’s anti-HIV effect. Overall, our studies concluded that LSM is a promising LPA to inhibit HIV-1 infection in myeloid cells in the RIOK1-dependent manner.

## Introduction

Despite the remarkable success of combination antiretroviral therapy (cART) in suppressing HIV-1 acute infection and plasma viremia in people living with HIV-1 (PLWH), a definitive cure remains elusive due to the persistence of transcriptionally silent, yet replication-competent HIV-1 proviruses integrated into the host genome, forming stable viral reservoirs(1, 2). These HIV-1 viral reservoirs are resistant to cART regimens and also capable of escaping immune surveillance, which induces the chronic immune dysfunctions and inflammation. Therefore, HIV-1 cure is actively pursued to disrupt HIV-1 persistent infection and/or HIV-1 induced immunopathogenesis.

While CD4+ T cells are considered as the major types of HIV-1 viral reservoirs, myeloid cells, including monocytes, microglia, and macrophages, are also susceptible to HIV-1 infection, which have recently gained the increasing attentions regarding their roles as long-lived viral reservoirs to support HIV-1 persistence(3–6). These cells are generally resistant to HIV-induced cytopathic effects. HIV-1 establishes the sustained infection and undergoes compartmentalized replication in tissue-resident macrophages, such as microglia at the central nervous system (CNS) and Kupffer cells in the liver, where antiretroviral drugs have the poor penetration and/or reduced immune clearance (7). Myeloid cells also exhibit the distinct mechanisms of HIV-1 latent establishment and maintenance. Earlier studies showed that HIV-1 transcription and latency regulations in myeloid cells differ substantially from those in CD4+ T cells, complicating their management and elimination(8).

Various therapeutic approaches have been tempted for curing HIV-1. Early efforts have been paid to develop the “shock and kill” approach for purging latent HIV-1 proviruses by using latency-reversing agents (LRAs), which is expected to induce a viral cytopathic effect that leads to clearance of HIV viral reservoirs and a sterilizing cure of HIV-1(9). Although certain progresses have been made, such approach is so far not successful to significantly reduce HIV-1 viral patients in clinical trials (10, 11). There still remains several safety concerns, especially the CNS complications induced by HIV-1 reactivation. On the contrary, the alternative “block and lock” approach has been developed with the goal to permanently silence HIV-1 proviruses, thus locking them in deep latent state and preventing episodic reactivation or viral “blips” to replenish viral reservoirs, which is expected to eventually reach a functional cure of HIV-1(12, 13). Several latency-promoting agents (LPAs) have been reported to hold a promise for blocking HIV-1 latent reactivation and delaying viral rebound with cART withdrawal, but most of these studies primarily focused on characterizing the impact of LPAs on HIV-1 viral reservoirs in CD4+ T cells. It is critical to develop LPAs that are also effective to target HIV-1 proviruses in myeloid cells, including both circulating monocytes and tissue-resident macrophages that contain stable HIV-1 viral reservoirs.

As an ideal LPA, it shall be safe, potent, and also capable of suppressing HIV-1 proviruses in multiple types of cells, including both CD4+ T cells and myeloid cells. We previously screened an FDA-approved drug library and identified levosimendan (LSM) as a promising LPA that blocks HIV-1 lytic reactivation in CD4+ T cells without obvious cytotoxicity (14). LSM is a calcium sensitizer and potassium channel opener approved for treatment of acutely decompensated severe chronic heart failure (ADHF) in Europe and other countries(15), but not approved in the United States. LSM is recognized for its effects on systemic and pulmonary hemodynamics and its capability to alleviate symptoms of acute heart failure (16). Due to its promising anti-HIV effects, we continued to evaluate the therapeutic potential of LSM as a novel LPA. In this study, we identified that LSM is able to block HIV-1 lytic infection and reactivation in myeloid cells beyond CD4+ T cells, supporting its broad applicability as an LPA. Based on a previous study indicating that the serine/threonine-protein kinase RIO1 (RIOK1) is a leading kinase target of LSM (17), we found that LSM suppresses HIV proviral gene expression in a RIOK1-dependent manner. Overall, our new findings confirm that LSM is a promising LPA to inhibit HIV infection and lytic reactivation in both myeloid cells and CD4+ T cells, which could be considered for “block and lock” HIV-1 cure strategies. Investigations of RIOK1 as a potential target of LSM also shed light in understanding the molecular mechanisms governing HIV latency.

## Results

### LSM inhibited HIV lytic reactivation in myeloid cell models of HIV-1 latency

We first determined the anti-HIV activities of LSM in HC69 cells that are derived from the immortalized C20 parental microglial cells. HC69 cells carry a latently infected, replication-defective HIV-1 NL4–3 viral construct that contains regulatory and structural genes (*Tat*, *Rev*, *Env*, *Vpu*, and *Nef)* with a d2EGFP reporter gene but lacks *Gag*, *Pol*, and certain accessory genes (*Vif*, *Vpr*). Our results showed that LSM treatment decreases the percentage of GFP-positive HC69 cells induced with TNFα (**Fig. 1A**). We further determined the IC_50_ value of LSM to inhibit HIV-1 lytic reactivation in HC69 cells (IC_50_ = 6.85µM) by calculating the relative percentage of GFP-positive cells (**Fig. 1B**). We also monitored the cell viability of LSM-treated HC69 cells and measured its LC_50_ (LC_50_ = 27.47 µM). At a high dose (25 µM), LSM resulted in the reduction of HIV-1 lytic reactivation by 97.3% while the cell viability still remained above 50%.

**Figure 1.**
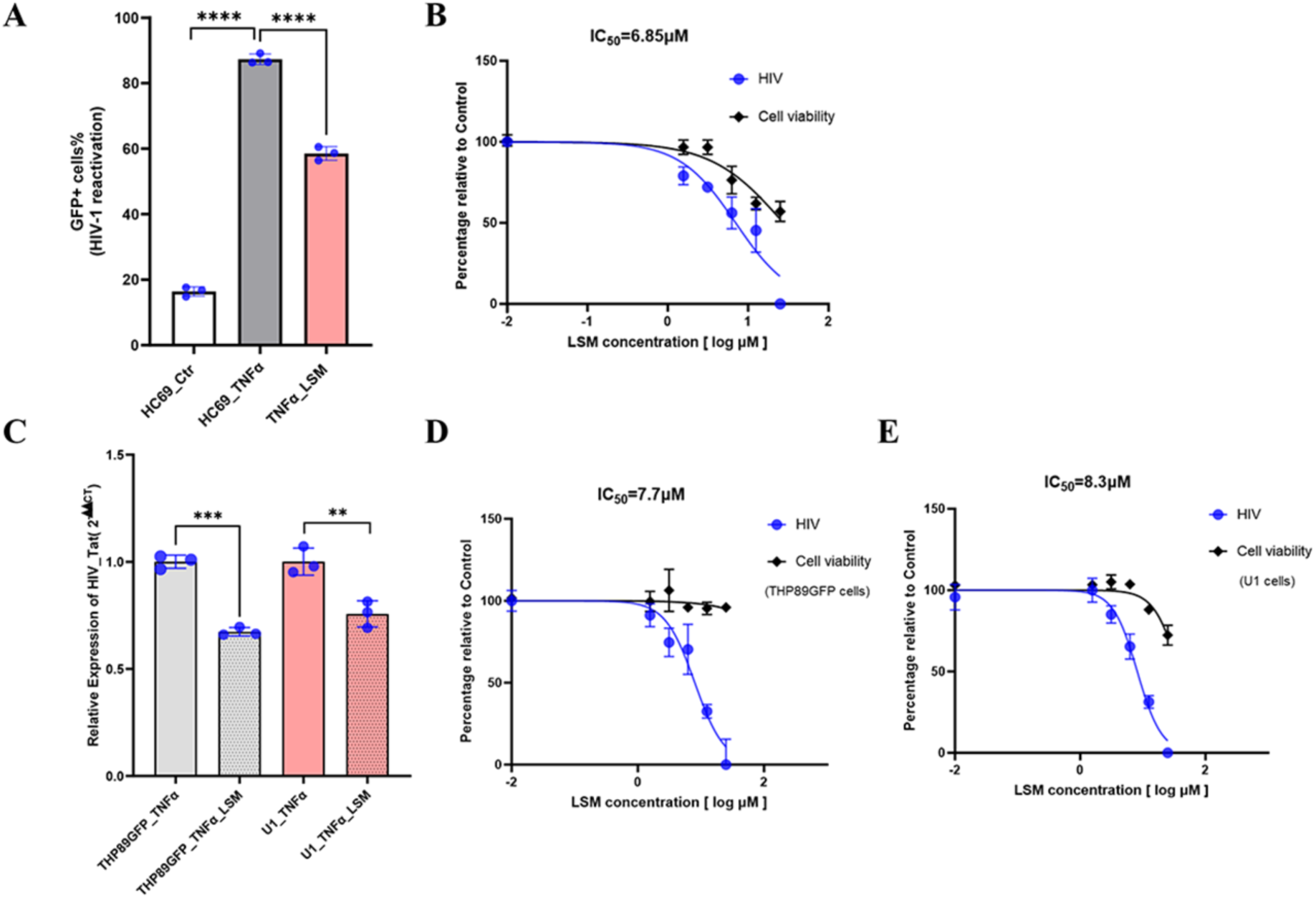
Levosimendan (LSM) inhibited HIV lytic reactivation in myeloid cell models of HIV-1 latency. **(A)** HC69 cells were treated with LSM (6.25 µM) for 24 hours and then co-treated with TNFα (10 ng/ml) for additional 48 hours. Cells were imaged using a Cytation 5 Cell Imaging Multi-Mode Reader for GFP expression, and the percentage of GFP-expressing cells was calculated by counting the number of GFP-positive cells relative to the total number of Hoechst-positive cells. (**B**) HC69 cells were titrated with LSM at a series of concentrations, and the percentage of GFP-positive cells was determined and plotted to determine the IC_50_ value. **(C)** THP89GFP and U1 cells were treated with LSM (6.25 µM) for 24 hrs and then co-treated with TNFα (10 ng/ml) for additional 48 hrs. The total RNAs were extracted and subjected to RT-qPCR analysis to determine the relative expression level of HIV-1 Tat. TNFα-induced cells without LSM treatment were used as a reference. **(D, E)** The THP89GFP **(D)** or U1 **(E)** cells were per-treated with LSM at a series of concentrations (0-25nM) for 24 hours and then co-treated with TNFα (10 ng/ml) for an additional 48 hours. Total RNAs were extracted for RT-qPCR analysis of HIV-1 Tat to determine the IC_50_ value. The cell viability was measured using CellTiter-Glo assay. TNFα-induced cells without LSM treatment were used as a reference. Results were presented as mean ± SD from three independent experiments and analyzed by a two-tailed Student’s t-test (**p < 0.01, ***p < 0.001, ****p <0.0001).

We further evaluated the anti-HIV potential of LSM in two monocytic cell lines of HIV-1 latency, THP89GFP and U1 cells. U1 cells are a chronically HIV-1-infected monocytic cell line, whereas THP89GFP cells are an HIV-1-infected monocytic latency reporter cell line. Both monocytic cells were treated with TNFα to induce HIV-1 lytic reactivation. Our results showed that LSM treatment decreases the TNFα-induced HIV viral gene expression in both of these cells (**Fig. 1C**). We also determined the IC_50_ value of LSM to inhibit HIV-1 lytic reactivation in these cells (THP89GFP: IC_50_ = 7.70µM; U1: IC_50_ = 8.30µM), which was in the similar range as HC69 cells (**Fig. 1D**, **1E**). We also measured the cell viability of these cells treated with LSM in parallel (THP89GFP: LC_50_ = 121.50 µM; U1: LC_50_ = 39.98 µM). It appeared that these monocytic cell lines (THP89GFP, U1) can tolerate LSM better than microglia (HC69). Overall, we demonstrated that LSM is capable of inhibiting HIV-1 lytic reactivation in several myeloid cell lines with tolerable cytotoxicity.

### LSM suppressed HIV-1 proviral transcription in human induced pluripotent stem cell (iPSC)-derived microglia

Beyond the use of microglia cell lines (HC69), we also included the HIV-infected iPSC-derived microglia (iMG) to determine the anti-HIV activities of LSM. iPSCs were first differentiated into hematopoietic progenitor cells (HPCs), which were subsequently differentiated into microglial cells. We confirmed that iMG express microglia/macrophage markers including ionized calcium-binding adapter molecule 1 (IBA1) and C-X3-C Motif Chemokine Receptor1 (CX3CR1) (18, 19) by protein immunofluorescence (**Fig. 2A**) and immunoblotting (**Fig. 2B**) analyses. These iMG were then pre-infected with the wild-type HIV-1Bal strain to allow the formation of proviruses, followed by LSM treatment. Our results showed that LSM suppresses HIV-1 *Tat* and *Gag* viral gene expression (**Fig. 2C, 2D)** as well as HIV-induced IL-1β expression (**Fig. 2E**) by RT-qPCR. This evidence supported that LSM is useful to suppress both HIV proviral transcription and HIV-associated inflammatory responses in microglia using the more advanced cell model of human microglia iMG that could overcome the limitations of animal models and inaccessible human samples.

**Figure 2.**
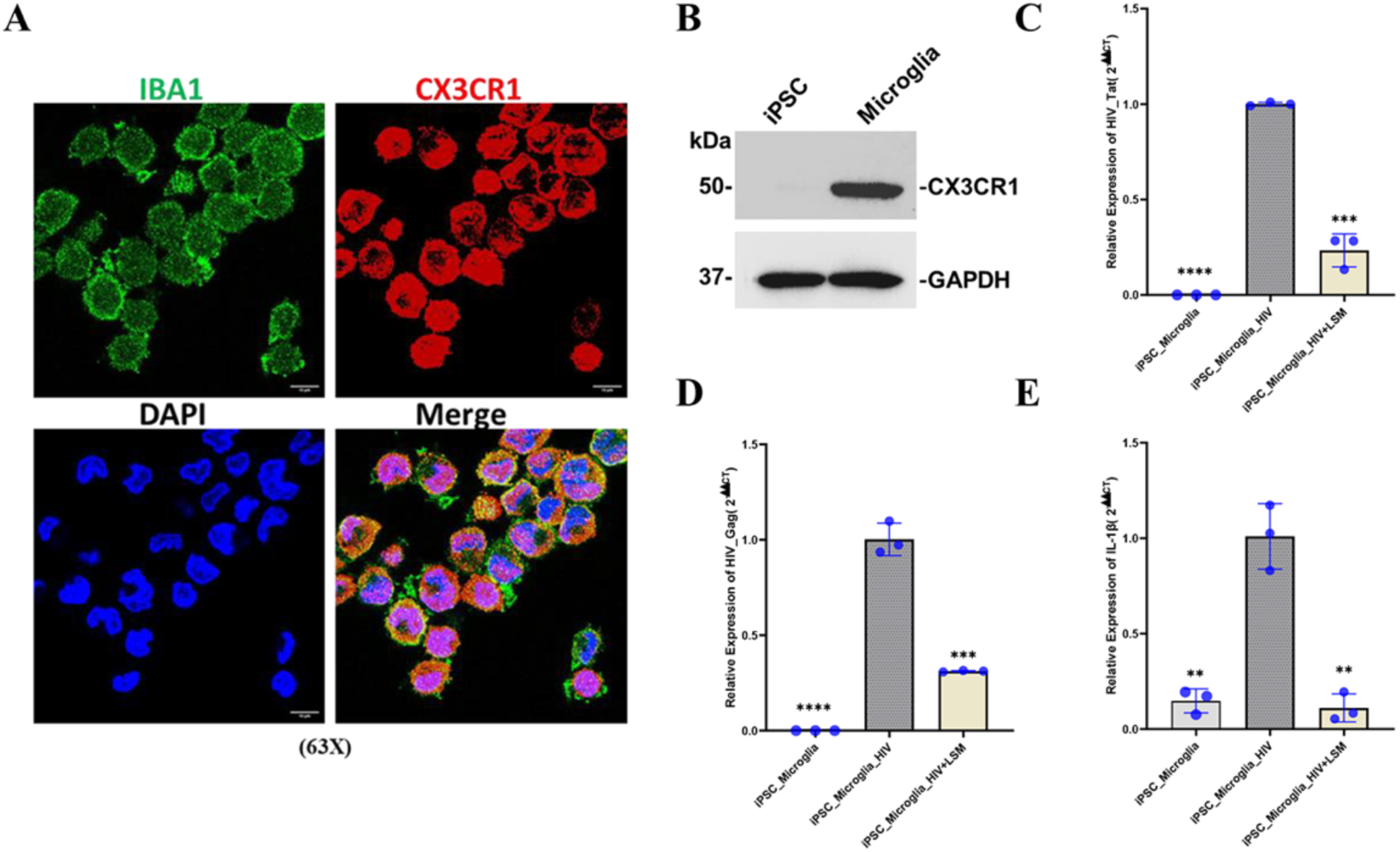
LSM suppressed HIV-1 proviral transcription in iPSC-derived microglia (iMG). **(A)** The microglia markers of generated iMG were verified by protein immunofluorescence analysis, including IBA1 (green) and CX3CR1 (red). The nuclei were stained with DAPI (blue). Scale bar: 10 μm. **(B)** Total proteins were extracted from iMG or original iPSC cells and analyzed by protein immunoblotting of microglia marker CX3CR1. GAPDH was used as a loading control. **(C-E)** iMG were infected with HIV-1 (MOI=1) for 5 days. Cells were then treated with LSM (6.25 µM) or mock treated for 48 hrs. Total RNAs were extracted for RT-qPCR analysis of HIV-1 Tat **(C)**, Gag **(D),** or IL-1β (**E**). HIV-infected cells without LSM treatment were used as a reference. Results were presented as mean ± SD from three independent experiments and analyzed by a two-tailed Student’s t-test (**p < 0.01, ***p < 0.001, ****p <0.0001).

### LSM suppressed HIV-1 proviral transcription in human primary stellate macrophages (Kupffer cells, KC) of liver

Our characterization of LSM further used the more physiologically relevant human primary macrophages, including human kupffer cells – stellate macrophages in liver. Using a human HeLa cell line (MAGI) as a reference, we confirmed that KC in our studies indeed express the macrophage-specific markers (CD45, CD163) by protein immunoblotting (**Fig. 3A**). Likewise, KC were then pre-infected with the wild-type HIV-1Bal strain to allow the formation of proviruses, followed by LSM treatment. Our results showed that LSM treatment suppresses HIV-1 proviral gene expression by protein immunoblotting of HIV-1 p24 (**Fig. 3B**) as well as RT-qPCR of HIV-1 *Tat* and *Gag* viral transcripts (**Fig. 3C, 3D**). These assays confirmed the anti-HIV potency of LSM by using human primary macrophages in liver, underscoring its potential to targeting HIV proviral transcription and lytic reactivation in tissue-resident macrophages (TRMs).

**Figure 3.**
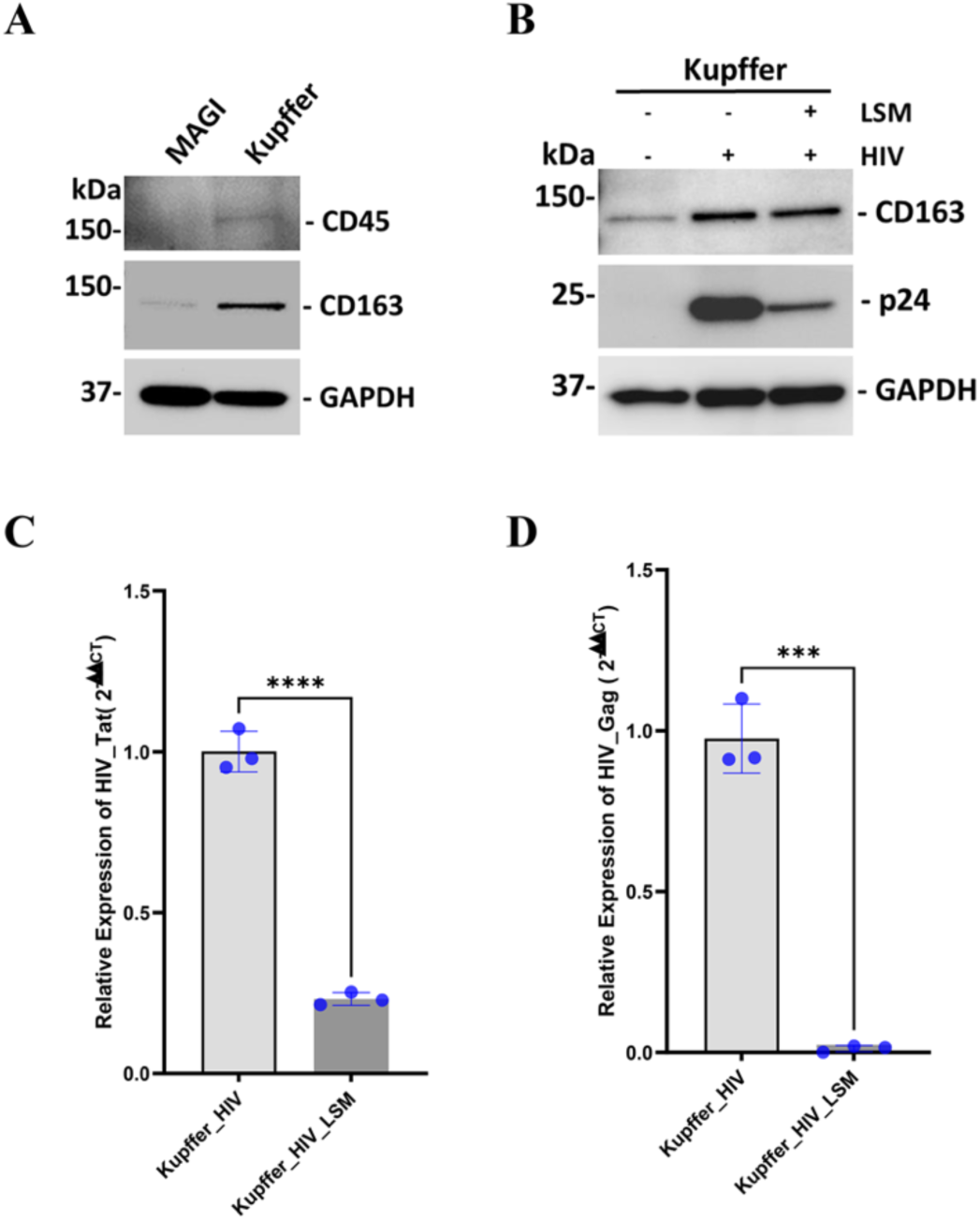
LSM suppressed HIV-1 proviral transcription in human primary liver macrophages. **(A)** The markers of Kupffer cells were verified by protein immunoblotting, including CD45 and CD163. GAPDH was used as a loading control. **(B)** The Kupffer cells were infected with HIV-1 (MOI=1) for 48 hours, and then treated with LSM (6.25 µM) for 48 hours. The cells were harvested and analyzed by protein immunoblotting for HIV p24, CD163, and GAPDH. **(C, D)** The Kupffer cells were infected with HIV-1 and treated with LSM as in (B). Total RNAs were extracted and analyzed by RT-qPCR to measure expression of HIV Tat **(C)** or Gag **(D)**. Results were presented as mean ± SD from three independent experiments and analyzed by a two-tailed Student’s t-test (***p < 0.001, ****p <0.0001).

### LSM blocked HIV-1 infection in human primary monocyte-derived macrophages (MDMs)

Macrophages derived from human primary monocytes were used as another model of TRMs to determine the anti-HIV potency of LSM. Purified human peripheral blood monocytes were differentiated to macrophages during the *in vitro* cultures. We tested the first scenario that MDMs were pre-treated with LSM prior to challenge of the wild-type HIV-1 Bal strain. Total RNAs were extracted and analyzed by RT-qPCR analysis. Our results showed that LSM blocks HIV acute infection through measurement of *Gag* viral transcript at both low and high doses (5 µM, **Fig. 4A**; 20 µM, **Fig. 4B**). In addition, LSM elicited almost no cytotoxicity in MDMs measured by the MTT assay across all tested doses (5-100 µM, **Fig. S1A**). LSM treatment generated no obvious impact on gene expressions of HIV-1 receptors either (CD4, **Fig. S1B**; CCR5, **S1C**).

**Figure 4.**
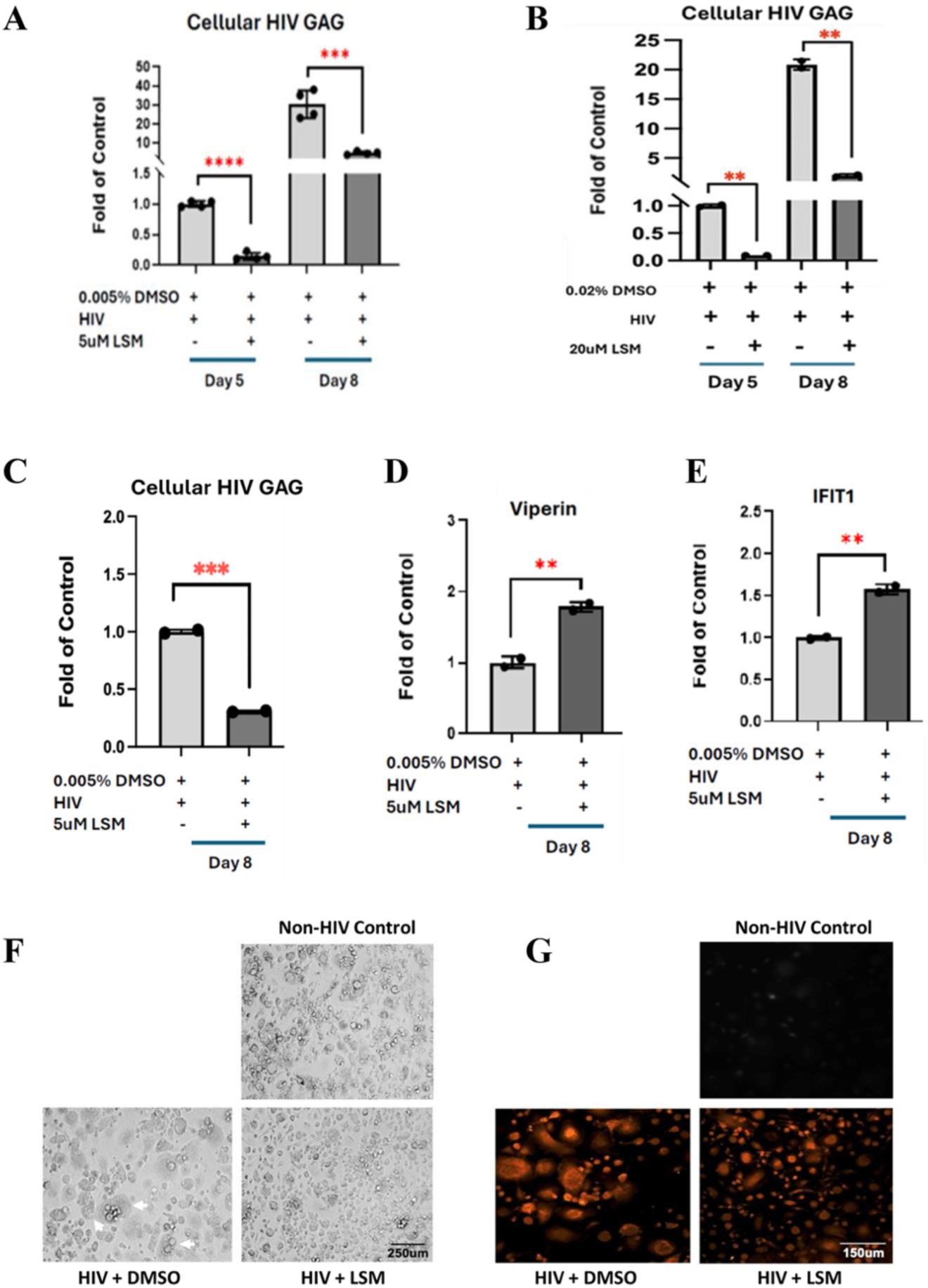
LSM blocked HIV-1 infection in human primary monocyte-derived macrophages. **(A, B)** Human MDMs were pretreated with LSM at 5 uM (A) or 20 uM (B) for 24 hours and then infected with HIV-1 Bal strain. Cells were collected at 5 or 8 days post of infection (dpi), followed by RNA extractions. Relative expression level of HIV Gag was analyzed by RT-qPCR. HIV-infected cells without LSM treatment were used as a reference. **(C-E)** Human MDMs were infected with HIV-1 Bal strain for 4 days and then treated with LSM (5 µM) for additional 8 days. Cells were collected for RNA extractions. Relative expression level of HIV Gag **(C)**, Viperin **(D),** or IFIT1 **(E)** was analyzed by RT-qPCR. HIV-infected cells without LSM treatment were used as a reference. Results were presented as mean ± SD from three independent experiments and analyzed by a two-tailed Student’s t-test (**p < 0.01, ***p < 0.001, ****p <0.0001). **(F)** Human MDMs were pretreated with LSM (5 uM) for 24 hours and then infected with HIV-1 Bal strain. At 8 dpi, cells were visualized for HIV-induced syncytia formation. Scale bar: 250 μm. (**G**) Human MDMs were infected with HIV-1 Bal strain for 4 days and then treated with LSM (20 µM) for additional 8 days. Cells were collected and analyzed by protein immunofluorescence for HIV p24 (red). Scale bar, 150 μm.

Alternatively, we tested the second scenario that MDMs were infected with the wild-type HIV-1 Bal strain prior to LSM treatment. Our results showed that LSM treatment consistently suppresses HIV-1 viral gene expression this scenario (**Fig. 4C**). Intriguingly, we noticed that LSM treatment also leads to the significant upregulation of certain antiviral innate immune genes (Viperin, **Fig. 4D**; IFIT1, **Fig. 4E**), which indicated that LSM might inhibit HIV-1 infection through both the direct impact on suppressing HIV proviral transcription and the indirect impact on activating anti-HIV innate immune responses. Furthermore, we confirmed that at both scenarios LSM potently blocked HIV-induced syncytia formation in human primary MDMs by bright-field microscopy or HIV p24 immunofluorescence microscopy **(Fig. 4F, 4G)**. Therefore, we showcased that LSM blocks HIV-1 infection in human primary MDMs used as another physiologically relevant model of human TRMs.

### LSM blocked HIV-1 infection in a RIOK1-dependent manner

Although we demonstrated that LSM possesses anti-HIV activities in both CD4+ T and myeloid cells, its mode of action remains unclear. LSM is mainly known to target the Ca^2+^-bound form of cardiac troponin C to stabilize the troponin complex, but an earlier report unraveled that LSM acts as a multi-target agent with significant off-target kinase inhibition, particularly against RIOK1, through computational screening (3D-REMAP) and experimental validations(17). LSM inhibited RIOK1 via the interaction with its ATP-binding pocket, leading to anticancer effects (20–22). RIOK1 belongs to the RIO (right open reading frame) family of atypical protein kinases/ATPases that also include RIOK2 and RIOK3, while RIOK1 is the mostly studied one. RIOK1 plays a role in ribosome biogenesis and cell cycle progression by supporting the release of biosynthetic factors during the maturation of pre-40S small ribosomal subunit (22, 23). RIOK1 also modulates p53 signaling by regulating its stability and activity (24). We thus determined whether LSM blocks HIV-1 infection in a RIOK1-dependent manner.

We first used MAGI cells for such studies. MAGI cells were pre-infected with HIV-1 IIIB strain followed by LSM treatment, which led to the decrease of HIV viral gene expression measured by RT-qPCR of *Tat* viral transcript (**Fig. S2A**) and immunoblotting of HIV p24 protein (**Fig. S2B**). To determine the impact of RIOK1, MAGI cells were transiently transfected with a vector expressing V5-tagged RIOK1 or control vector, followed by HIV-1 infection and LSM treatment. Our results showed that overexpression of RIOK1 enhances HIV infection while LSM treatment abolishes such effects (**Fig. 5A**, **Fig. S2C**).

**Figure 5.**
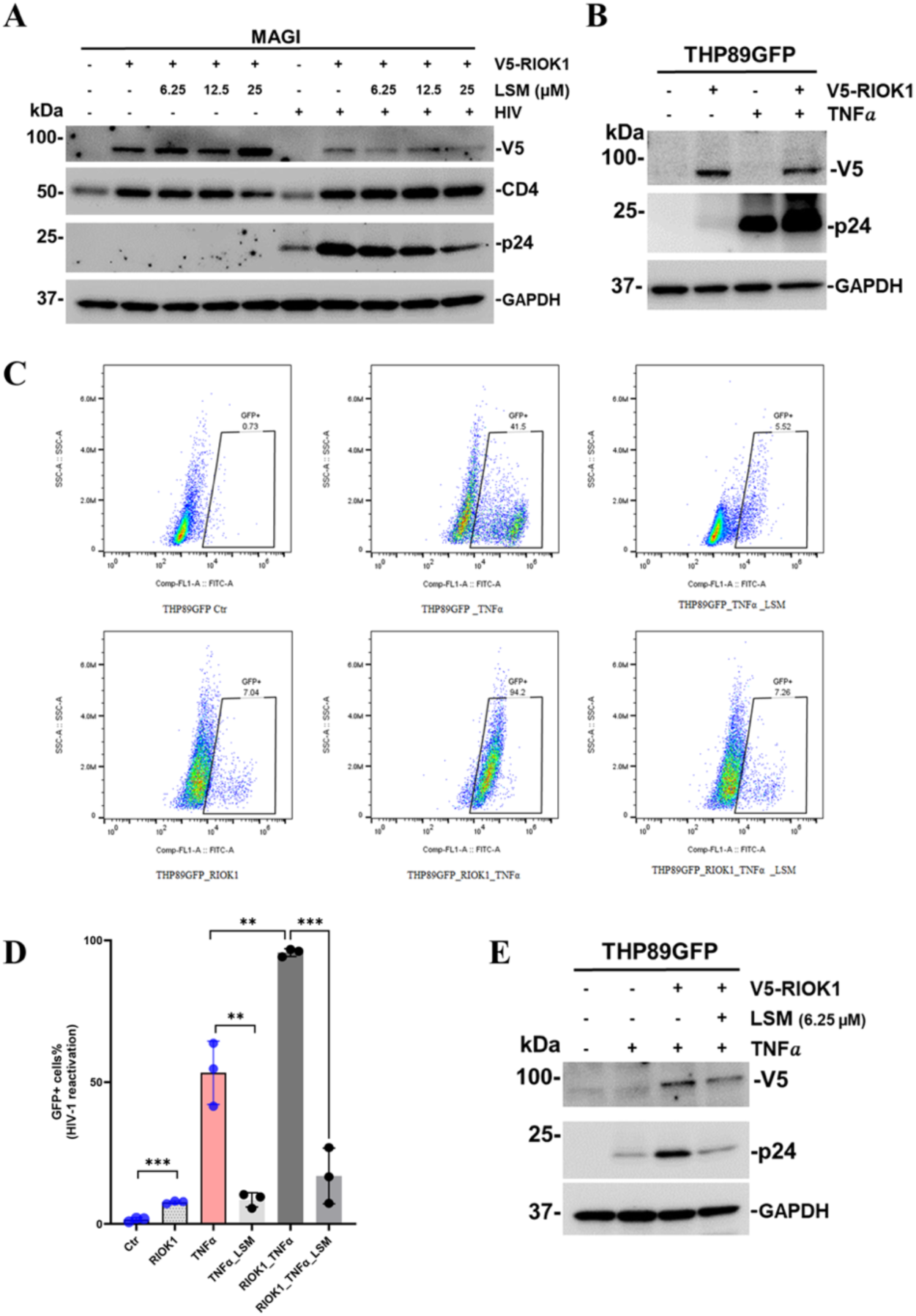
LSM blocked HIV-1 infection in a RIOK1-dependent manner. **(A)** MAGI cells were transiently transfected with the vector expressing V5-tagged RIOK1 or the empty vector as a control for 48 hours, followed by HIV-1 infection (MOI=1) for 48 hours. Cells were further treated with LSM at the indicated concentrations for additional 48 hours. Total proteins were extracted and analyzed by protein immunoblotting for HIV p24, CD4, V5, and GAPDH. **(B)** THP89GFP cells were transiently transfected with the vector expressing V5-tagged RIOK1 or the empty vector as a control for 48 hours, followed by treatment of TNFα (10 ng/ml) for 48 hrs. The cells were harvested and lysed for immunoblotting analysis using antibodies against HIVp24, V5 and GAPDH. GAPDH was served as a loading control. **(C-E)** THP89GFP cells were transiently transfected with the vector expressing V5-tagged RIOK1 or the empty vector as a control for 48 hours, followed by treatment of TNFα (10 ng/ml) with or without LSM (6.25 µM) for additional 48 hours. (**C**). Cells were fixed and analyzed by flow cytometry at the SSC-A and FITC-A (GFP) channels. The plots of one individual experiment were illustrated. (**D**). The percentage of GFP-positive cells indicating the rate of HIV lytic reactivation was quantified. Results were presented as mean ± SD from three independent experiments and analyzed by a two-tailed Student’s t-test (***p < 0.001). (**E**). Proteins in above cells were extracted and analyzed by protein immunoblotting for HIV p24, V5, and GAPDH. GAPDH was served as a loading control.

We further confirmed these findings in THP89GFP cells. Overexpression of RIOK1 further enhanced TNFα-induced HIV-1 lytic reactivation in these cells measured by immunoblotting of HIV-1 p24 protein (**Fig. 5B**). The above cells were treated with or without LSM, followed by flow cytometry analysis of HIV-1 lytic reactivation. LSM indeed abolished the impact of RIOK1 overexpression on enhancing TNFα-induced HIV-1 lytic reactivation in THP89GFP cells measured by percentage of GFP-expressing cell population (**Fig. 5C, 5D**). Such effects of LSM in a RIOK1-dependent manner were also verified by immunoblotting of HIV-1 p24 protein (**Fig. 5E**). Overall, these results suggested that LSM likely acts through targeting and inhibiting RIOK1 functions to block HIV-1 infection.

## Discussion

Myeloid cells, including monocytes and tissue-resident macrophages/microglia, are pivotal in HIV-1 persistence, acting as long-lived viral reservoirs despite cART(25). These cells, particularly macrophages/microglia, harbor HIV proviral DNAs, undergo low-level viral replications, and produce replication-competent viruses over the long periods. A significant feature of HIV-1 infected macrophages/microglia is the resistance to viral-induced cytopathic effects that generally kill CD4+ T cells, making them a significant barrier to reach a HIV sterilizing cure by considering “shock and kill” strategy that primarily relies on HIV-1 induced cell death(26, 27). On the contrary, the “block and lock” approach designed to permanently silence the HIV-1 genome (“deep latency”) might be applied to long-lived myeloid cellular reservoirs to reach a HIV-1 functional cure(28).

We previously identified that LSM is potent to block HIV-1 lytic reactivation in CD4+ T cells, and our new results demonstrated that LSM is also effective to block HIV infection and lytic reactivation in myeloid cells as well. Furthermore, it remains to be determined how LSM suppresses HIV-1 proviral transcription despite of our early efforts showing that the anti-HIV effects of LSM is not through its interference with either HIV-1 Tat protein expression or interaction of Tat with the Positive Transcription Elongation Factor b (P-TEFb) complex(14). Nevertheless, our new findings suggested that LSM might inhibit HIV in a RIOK1-depdentn manner, which would provide new clues to further dissect the molecular mechanisms underlining LSM’s anti-HIV activities. On the other hand, LSM could be used as a chemical probe of RIOK1 to understand its previously unappreciated biological functions to promote HIV infection in future.

Although it is not approved by the FDA in the United States, LSM is primarily used throughout the European Union and Latin America as an intravenous inotropic agent for the short-term treatment of acute decompensated heart failure (15, 16). LSM increases calcium sensitization in primary guinea pig myocytes at the range of 1 nM to 10 µM, while the relatively high concentrations (30-100 nM) are needed for the cytoprotective effects against apoptosis in primary rat cardiac fibroblasts(29). Our results showed that LSM modestly reduces the cell viability of HC69 and U1 cells only at the high concentrations (**Fig. 1B, 1E**), while LSM had almost no impact on the cell viability of THP89GFP cells at all tested concentrations (**Fig. 1D**). Likewise, we showed that human primary MDMs tolerate LSM well with no obvious cytotoxicity even at the high doses up to 100 µM (**Fig. S1A**). For most of our LSM studies *in vitro* using various myeloid cell models, we selected the doses (around 5 to 20 µM) that cause no or modest cytotoxicity yet potently inhibit HIV-1 (**Fig. 2-4**). Given LSM has already been used in humans, it is likely safe to further move forward to initiate animal and/or pre-clinical studies to further evaluate its anti-HIV activities.

Alongside its direct effects on silencing HIV-1 proviral transcription, LSM appeared to regulate host immune and inflammatory responses as well. Interestingly, LSM treatment led to the upregulation of antiviral ISGs, such as Viperin and IFIT1 (**Fig. 4D, 4E**), both of which have been reported to inhibit HIV-1 viral production in human macrophages (30, 31). LSM may elicit the anti-HIV activities through direct and indirect mechanisms. Furthermore, we showed that LSM blocks HIV-1 induced upregulation of proinflammatory cytokine IL-1β (**Fig. 2E**). Indeed, early studies have demonstrated that beyond its primary inodilator function LSM exhibits the strong anti-inflammatory properties by reducing pro-inflammatory cytokines (IL-6, TNF-a, IL-1β) while inhibiting NF-kB activation (32, 33). Thus, we believe that LSM is a promising therapeutic candidate to treat both HIV-1 infection and associated chronic inflammation due to its multivalent functions to target HIV-1 as well as host immune and inflammatory gene expressions. However, a more thorough characterization of LSM’s impact on the entire host transcriptome in HIV-infected myeloid cells is needed while the participating cellular signaling factors and pathways remain to be further investigated.

One of key findings from this study is that the anti-HIV activities of LSM likely depends on RIOK1 (**Fig. 5**). RIOK1 is an atypical serine/threonine kinase involved in ribosome biogenesis and cell survival(34). RIOK1 has not been previously reported to regulate viral infections. Known host regulators of HIV-1 transcription include NF-kB, Sp1, and P-TEFb/CDK9 complex, but RIOK1 has not been implicated in any parts of these pathways in previous publications. To the best of our knowledge, this is the first time to show that RIOK1 might be required for active HIV-1 proviral transcription. RIO kinases have unique ATP-binding features, and the single-copy presence per subfamily enables the development of highly specific inhibitors (35). One early study indicated that RIOK1 is a major ATP-competitive target of LSM, showcasing its multi-target kinase inhibitory activities (17). Our previous work also demonstrated that LSM inhibits HIV-1 lytic reactivation likely through modulation of PI3K pathway (14). As PI3K functions upstream of AKT, PI3K-mediated PIP3 production leads to AKT activation. RIOK1 indeed phosphorylates and activates AKT as well. It has been shown that knockdown of RIOK1 disrupts ribosome biogenesis and AKT signaling pathway (22, 36). Thus, LSM likely targets PI3K-RIOK1-AKT signaling axis, which shall be further studied in future. Interestingly, overexpression of RIOK1 synergized with TNF*a* stimulation to reactivate latent HIV-1, indicating that LSM could be combined with other HIV-1 LPAs to maximize the anti-HIV effects on preventing HIV-1 lytic reactivation and thus reinforcing its viral latency, leading to a functional cure.

## Materials and Methods

### Cell lines and cultures

HC69 cells were cultured in BrainPhys™ Neuronal Medium (STEMCELL Technologies) supplemented with N-2 supplement (Thermo Fisher Scientific), L-Glutamine (2.6 mM), 10% Fetal bovine serum (FBS), Normocin (100 µg/ml), penicillin (100 U/ml) and streptomycin (100 μg/ml). MAGI cells were cultured in Dulbecco’s Modified Eagle Medium (DMEM) supplemented with 10% FBS, penicillin (100 U/ml) and streptomycin (100 μg/ml). THP89GFP and U1 cells were maintained in RPMI 1640 medium supplemented with 10% FBS, penicillin (100 U/ml) and streptomycin (100 μg/ml). Human primary Kupffer cells (Cat# HLKC-200K, Lonza) were cultured in KuBM Kupffer cell basal medium supplemented with KuGM^TM^ SingleQuots (Cat# MKC-500SQ, Lonza). iPSC (Cat# SCTi003-A, STEMCELL) were cultured in mTeSR^TM^ Plus (Cat# 100-0274, STEMCELL) supplemented with mTeSRTM Plus 5X supplement (Cat# 100-0275, STEMCELL). All cells were cultured in a 5% CO_2_ incubator at 37 °C. Purified human peripheral blood monocytes were obtained from the Human Immunology Core at the University of Pennsylvania. The monocytes were cultured in 96-well or 48-well plates (Corning CELLBIND Surface) at 37°C with 5% CO_2_ in DMEM containing 10% FBS, 2 mM L-glutamine, 50 units/ml penicillin and 50 μg/ml streptomycin. Monocytes differentiated to macrophages after *in vitro* cultured for 7 days. We used the 7-day-cultured macrophages for experiments of this study.

### Viruses and Plasmids

HIV-1 IIIB and BaL wild-type strains were kindly provided by NIH AIDS reagent program. The coding sequence of human RIOK1 lacking a stop codon was cloned into the pLVX-M-Puro vector (Gene Universal Inc.) by using XhoI and ApaI restriction sites, generating a construct that expresses the C-terminal V5-tagged RIOK1. The plasmid expressing V5-tagged RIOK1 was amplified in *E.coli*, and purified using the QIAGEN Plasmid plus Midi kit (Cat#12945, QIAGEN). For adherent cells, they were transfected with the plasmid at the 80–90% confluence by using the FuGENE6 transfection reagent (Cat# E2691 Promage). For suspension cells such as THP89GFP, SusFexin (TS316, LAMA) was used for plasmid transfection according to the manufacturer’s instructions.

### Compounds and antibodies

Levosimendan (Cat# L5545) was purchased from Sigma. Recombinant human TNF*a* (Cat# BD554618) was purchased from BD Bioscience. HIV-1 p24 monoclonal antibody (IgG) was produced from the hybridoma cell line (NIH AIDS reagent program). The CD163 antibody (Cat# 16646-1-AP) was purchased from Proteintech Group. The CD45 antibody (Cat# MAB14302) was purchased from R&D Systems. Antibodies against IBA1 (Cat# MA5-27726), CX3CR1 (Cat# 702321), and V5 (Cat# 460705) were purchased from Invitrogen. The CD4 antibody (Cat# 93518S) was purchased from Cell Signaling Technology.

### Hematopoietic differentiation

According to the previous studies (25, 26), iPSCs were differentiated into hematopoietic progenitor cells (HPCs) by using the STEMdiff^TM^ Hematopoietic Kit (Cat# 05310, STEMCELL). Cells were maintained in mTeSRTM Plus medium on hESC-qualified Matrigel-coated 6-well plates. Differentiation was initiated by replacing mTeSRTM Plus with Hematopoietic Basal Medium A containing supplement A (1:200) at Day 1, followed by a half-medium change on Day 2 and replacement with Hematopoietic Basal Medium B containing supplement B (1:200) on Day 3. Half-medium changes were performed on Day 5, 7, 9, 10, and 12. The non-adherent, supernatant cells were collected on Day 12, and centrifuged at 300×g for 5 min. The cell pellets were collected and subjected to the downstream microglia differentiation.

### Microglia differentiation and maturation

The collected HPCs were further differentiated into iPSC-derived Microglia (iMGs) by culturing at a density of 2×10^4^ cells/cm^2^ on Matrigel-coated 6 well plates containing 2 ml STEMdiff^TM^ Microglia Differentiation Medium (Cat# 100-0019, STEMCELL). Cells were maintained at 37 °C with 5% CO_2_ and fed every other day by adding fresh medium (50% of the initial volume) without removing existing medium. On Day 12, the cells were suspended and collected, which were centrifuged at 300×g for 5 min. The cell pellets were resuspended in the remaining 1 ml of supernatant. The resuspended cells were then transferred to a new Matrigel-coated 6 well plates containing 1 ml fresh STEMdiff^TM^ Microglia Differentiation Medium and incubated at 37 °C with 5% CO_2_. Cells were fed every other day for an additional 12 days by adding fresh medium equal to half of the initial volume without removing the existing medium. On Day 24, cells were collected and centrifuged at 300×g for 5 min, and the resulting cell pellets were resuspended in the remaining 1 ml of supernatant. The resuspended cells were transferred to the freshly prepared Matrigel-coated 6 well plates containing 1 ml of STEMdiff^TM^ Microglia Maturation Medium (Cat#100-0020, STEMCELL) and maintained at 37 °C in a humidified incubator with 5% CO_2_. Cells were fed every other day for another 10 days by supplementing cultures with fresh maturation medium equal to 50% of the initial volume, without removal of existing medium. iPSC-derived Microglia (iMGs) were considered mature at Day 34 and subjected to various downstream studies.

### Protein immunoblotting

Cell pellets were lysed in RIPA lysis buffer (Cat# 20-188, Millipore Sigma) containing protease and phosphatase inhibitors. The lysates were then sonicated and centrifuged, and protein concentration in the supernatant was quantified by using the BCA assay kit (Cat #23227, Thermo Scientific). The protein samples were boiled in the SDS loading buffer and the equal amount of cell lysate (20 μg) was loaded into Novex™ WedgeWell™ 4–20% SDS-PAGE gels for protein separation, which was then transferred to a PVDF membrane. The membranes were blocked by 5% nonfat milk in TBST buffer (20 mM Tris-HCl, 150 mM NaCl, and 0.1% Tween 20) and incubated with the primary antibodies at 4 °C overnight, followed by the incubation with secondary antibodies at room temperature (RT) for 1 hour. Protein bands were visualized by using the Clarity Max ECL substrate (Cat# 1705062, Bio-Rad).

### Cell viability assays

Cell viability was mostly assessed by using the CellTiter-Glo® Luminescent cell viability kit (Cat#G7572, Promega). For MDMs, their cell viability was measured by MTT [3-(4,5-Dimethylthiazol-2-yl)-2,5-Diphenyltetrazolium Bromide].

### Protein immunofluorescence assay (IFA)

On Day 34 post of cell differentiation, iMGs were seeded onto glass coverslips coated with Poly-L-lysine and allowed to adhere for 24 hours prior to fixation. Cells were washed with cold PBS and fixed with 4% paraformaldehyde (PFA) for 20 min at RT. The cells were permeabilized by incubating with PBS containing 0.2% Triton X-100 for 15 minutes at RT. Cells were then washed and blocked, followed by incubating with the primary antibodies overnight at 4°C. Cells were then incubated with Alexa Fluor-488 or 568-conjugated secondary antibodies for 1 hour at RT, mounted with a DAPI-containing mounting medium, and imaged via a confocal microscope. GFP expression in HC69 was imaged as previously described (14). Briefly, cells were stained with Hoechst®33342 (Cat#62249, Thermo Fisher Scientific), and imaged using a Cytation 5 Cell Imaging Multi-Mode Reader. The percentage of GFP-positive cells was calculated by counting the number of GFP-expressing cells relative to the total number of Hoechst-positive cells.

### Flow cytometry

Cells were harvested, washed with PBS, and fixed with 4% paraformaldehyde. For flow cytometry analysis, cells were resuspended in PBS containing 2% BSA and analyzed by using a BD Accuri™ C6 Plus Flow Cytometer (BD Biosciences) at FSC, SSC-A, and FITC-A (GFP) channels. The percentage of fluorescence positive (GFP+) cells was measured and then quantified by using the FlowJo V10.8.1 software.

### Reverse-transcription (RT) coupled with real-time quantitative PCR (qPCR)

Total RNAs were extracted from the harvested cells by using the NucleoSpin RNA Extraction Kit (Cat#740955.250, MACHEREY-NAGEL). RNA samples were reverse transcribed into cDNAs by using the iScript^TM^ cDNA synthesis Kit (Cat# 1708891, Bio-Rad). Real-time qPCR was performed by using the iTaq Universal SYBR® Green Supermix (Cat#1725124, Bio-Rad). The qPCR reaction was carried out according to the manufacturer’s instructions under the conditions: 95℃ for 5 min, followed by 50 cycles of 95℃ for 15 sec, and 60℃ for 1 min. The relative expression levels of target genes were normalized to GAPDH by using the 2^−ΔΔCT^ method. The following qPCR primers were used in this study. HIV-1 Gag primers: Forward 5’-GAC GCT CTC GCA CCC ATC TC-3’, Reverse 5’-CTG AAG CGC GCA CGG CAA-3’; HIV-1 Tat primers: Forward 5’-GAAGCATCCAGGAAGTCAGC -3’, Reverse 5’-CTTCCTGCCATAGGAGATGC -3’; IL-1β primers: Forward 5’-CCACAGACCTTCCAGGAGAATG -3’, Reverse 5’-GTGCAGTTCAGTGATCGTACAGG-3’; GAPDH primers: Forward 5’-GTCTCCTCTGACTTCAACAGCG -3’, Reverse 5’-ACCACCCTGTTGCTGTAGCCAA -3’.

### Statistical Analysis

Data was presented as the mean ± SD of at least three independent biological experiments. Quantitative data were analyzed by using a two-tailed Student’s t-test. P values < 0.05 were considered statistically significant.

## Declaration of competing interest

The authors declare no competing interest.

## Acknowledgments

This study was supported by National Institute of Health (NIH) research grants R01DA059538, R01MH134402, R56AI157872 to J.Z.

## Supplementary Figures

**Figure S1.**
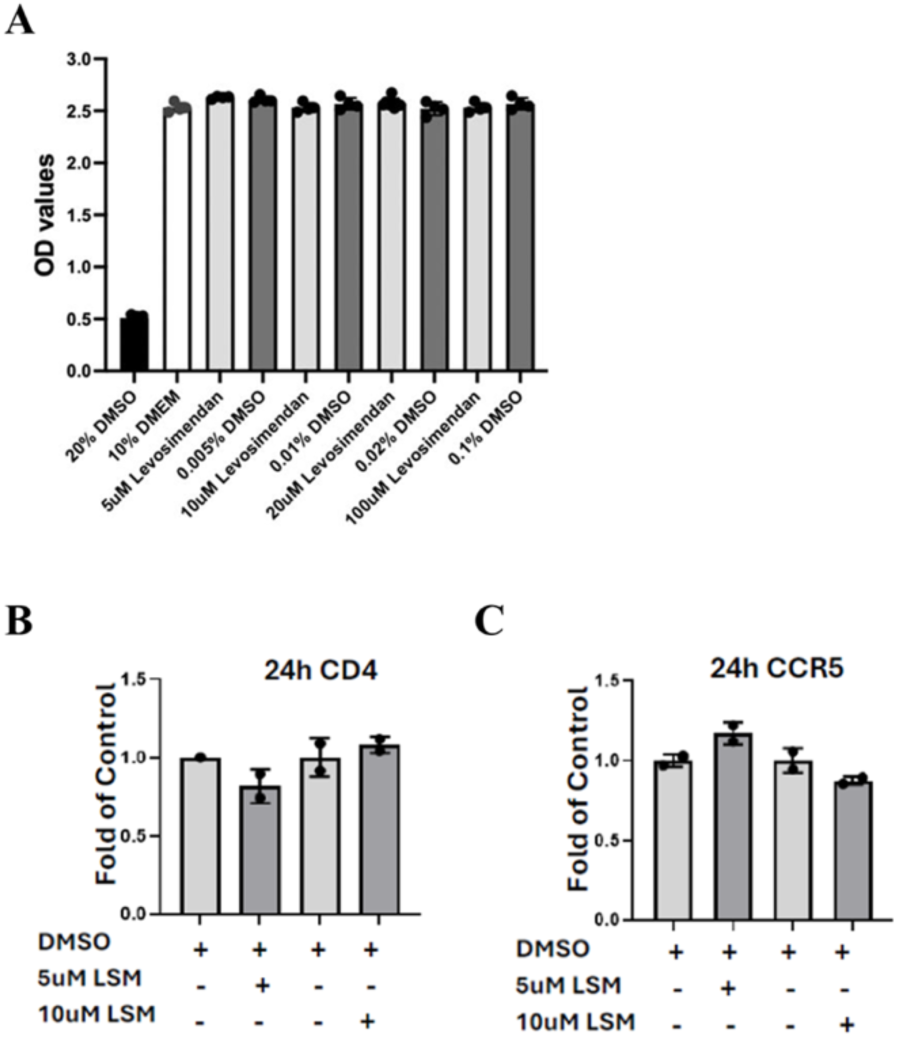
**(A)** Human MDMs were treated with LSM at a series of concentrations for 8 days. Cells were harvested and analyzed by the MTT assay to determine cell viability. **(B, C)** Human MDMs were treated with LSM (5µM or 10 µM) for 24 hrs. Total RNAs were extracted and analyzed by RT-qPCR to quantify the expression of CD4 (B) or and CCR5 (C).

**Figure S2.**
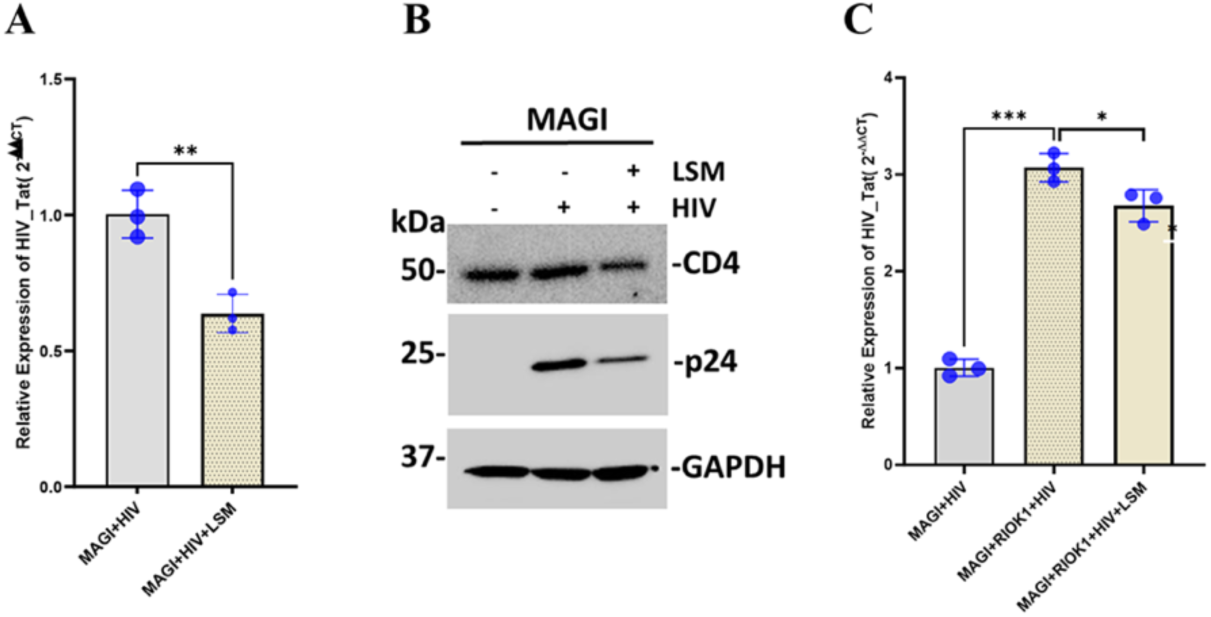
**(A)** MAGI cells were infected with HIV-1 (MOI=1) for 2 days and then treated with LSM (6.25µM) for additional 48 hrs. Total RNAs were extracted and analyzed by RT-qPCR to quantify the expression of HIV-1 Tat. **(B)** MAGI cells were infected with HIV-1 (MOI=1) for 48 hours and then treated with LSM (6.25 µM) for 48 hours. Total proteins were extracted and analyzed by protein immunoblotting for HIV p24 and CD4. GAPDH was used as a loading control. **(C)** MAGI cells were transiently transfected with the vector expressing V5-tagged RIOK1 or the empty vector as a control for 48 hours, followed by HIV-1 infection (MOI=1) for 48 hours. Cells were further treated with LSM (6.25 µM) for additional 48 hours. Total RNAs were extracted and analyzed by RT-qPCR to quantify the expression of HIV-1 Tat.

## Notes

### Competing Interest Statement

The authors have declared no competing interest.

